# Preferential transport of synaptic vesicles across neuronal branches is regulated by the levels of the anterograde motor UNC-104/KIF1A

**DOI:** 10.1101/2023.06.30.547240

**Authors:** Amruta Vasudevan, Neena Ratnakaran, Kausalya Murthy, Shikha Ahlawat, Sandhya P. Koushika

## Abstract

Asymmetric transport of cargo across axonal branches is a field of active research. Mechanisms contributing to preferential cargo transport along specific branches *in vivo* in wild type neurons are poorly understood. We find that anterograde synaptic vesicles preferentially enter the synaptic branch or pause at the branch point in *C. elegans* PLM neurons. The anterograde motor UNC-104/KIF1A regulates this vesicle behaviour at the branch point. Reduced levels of functional UNC-104 cause vesicles to predominantly pause at the branch point and lose their preference for turning into the synaptic branch. SAM- 4/Myrlysin, which aids in recruitment/activation of UNC-104 on synaptic vesicles, regulates vesicle behaviour at the branch point similar to UNC-104. Increasing the levels of UNC-104 increases the preference of vesicles to go straight towards the asynaptic end. This suggests that the neuron optimises UNC-104 levels on the cargo surface to maximise the fraction of vesicles entering the branch and minimise the fraction going to the asynaptic end.

## Introduction

Axonal branching is a complex, regulated phenomenon with crucial roles in establishing connectivity of neuronal circuits (Gallo, 2011). Axonal branches have distinct mechanisms of formation, leading to diverse morphologies with varied functions (Gibson & Ma, 2011). Axons can form i) terminal arbours with secondary, tertiary, and higher-order branches at sites of synapse formation, ii) bifurcations where two daughter processes innervate spatially disparate targets, or iii) collateral or interstitial branches, which are often formed far away from axon terminals and extend at either acute or right angles to the main process (Gibson & Ma, 2011, Gallo, 2011). Developmentally, there is an inherent asymmetry that is established in the formation of bifurcate and collateral branches. It is likely that these branches carry distinct molecular and biochemical signatures, as they respond to extracellular signals differently (Gibson & Ma, 2011), and can differ in the nature and extent of synaptic activity (Xue et al., 2014). Since branches formed from a single neuron can be heterogeneous in synapse formation and activity, it is reasonable to conclude that these branches may also differ from each other in the distribution of synaptic cargo. A recent study (Tymanskyj, Curran, and Ma, 2022) demonstrates that triggering of growth cone activity specifically at the end of one axonal branch of vertebrate neurons in culture leads to preferential transport of synaptic vesicles and LAMP1 vesicles into that branch. It is interesting to investigate whether axonal branches exhibit heterogeneous distribution of synaptic cargo in neurons *in vivo*, and the underlying mechanisms that bring this about.

Axonal branch points have been shown to contain curved microtubules (Yu et al., 1994, Gallo, 2011), intersecting microtubules (Letourneau, 1982), and microtubule distal ends (Yu et al., 1994). In addition, they contain stalled mitochondria (Spillane et al., 2013), patches of filamentous actin (Gallo, 2011, Chia et al., 2014), and clusters of stalled pre-SVs, which contribute to crowding (Sood et al., 2018). The complex cytoskeletal geometry coupled with the presence of the above-mentioned crowding agents make the branch point a crowded environment. In order to traverse the crowded environment of the branch point, the cargo– motor complex must navigate microtubule ends, obstructions such as stalled cargo, and cytoskeletal intersections. It has been demonstrated that vesicle behaviour at sites containing microtubule ends is determined by the density of laterally-aligned microtubules (Yogev et al., 2016, Gramlich et al., 2021) and the mechanics of the microtubule-dependent motors driving vesicle transport (Gramlich et al., 2021). Further, the behaviour of moving pre-SVs at sites of stalled cargo is reported to be affected by the levels of functional motor in the neuronal process (Sood et al., 2018). Upon reduction in levels of functional anterograde motor UNC- 104/KIF1A, anterogradely-moving pre-SVs predominantly stop at sites of stalled cargo, and exhibit a decrease in propensity to cross stationary clusters or emerge from them (Sood et al., 2018). Thus, the levels of motor in the neuronal process can be important determinants of pre-SV behaviour at neuronal branch points.

The motor–cargo complex can influence preferential transport by regulating the switching of microtubules at or near the branch point, from a microtubule going straight onto a track entering the branch. *In vitro* studies demonstrate that cargo behaviour at intersections is regulated by motor levels on the cargo surface and the forces generated by them (Bergman et al., 2018; Ross et al., 2008; Schroeder et al., 2010). These studies have significant implications for cargo transport at branch points *in vivo*. An *in vitro* study by Ross et.al., 2008 examined the behaviour of Kinesin-1 and the dynein-dynactin motor complex at microtubule intersections. It showed that beads transported by Kinesin-1 have a higher propensity to switch microtubule tracks at an intersection (∼70%) as opposed to pausing, travelling straight, or dissociating from the microtubules, while dynein-driven beads show a higher propensity to stop at intersections (Ross et al., 2008). A study conducted by Osunbayo et al., 2015, has observed that upon increasing the concentration of Kinesin-2 (KIF5A) on the surface of beads, they tended to pause longer at microtubule intersections. Schroeder et.al., (2010) has shown that switching of cargo between actin-microtubule intersections was governed by the number of engaged motors (kinesin v/s myosin) on the bead surface. In summary, *in vitro* studies show that cargo behaviour at branch points falls under three categories: 1) pausing at intersections, 2) switching tracks, and 3) continuing straight along the same track, and that motor levels on the cargo surface can influence these behaviours.

We use the *C. elegans* posterior lateral mechanosensory (PLM) neuron to investigate the behaviour of synaptic cargo at branch points *in vivo*, as it has a collateral branch forming chemical synapses (synaptic branch), while the main process does not (Fig. 1A). Synaptic vesicle precursors (pre-SVs) carry several important synaptic proteins on their membrane, but may not contain neurotransmitters in their lumen. They are transported in high numbers along the process and are required to turn into the synaptic branch for function. This is a good system to investigate whether pre-SVs exhibit a preference for the synaptic branch over the distal process without chemical synapses. We find that anterogradely-moving pre-SVs preferentially enter the synaptic branch as opposed to the distal process. This preferential transport is dependent on the levels of functional anterograde motor UNC-104/KIF1A in the neuronal process and likely on pre-SVs.

**Figure 1:**
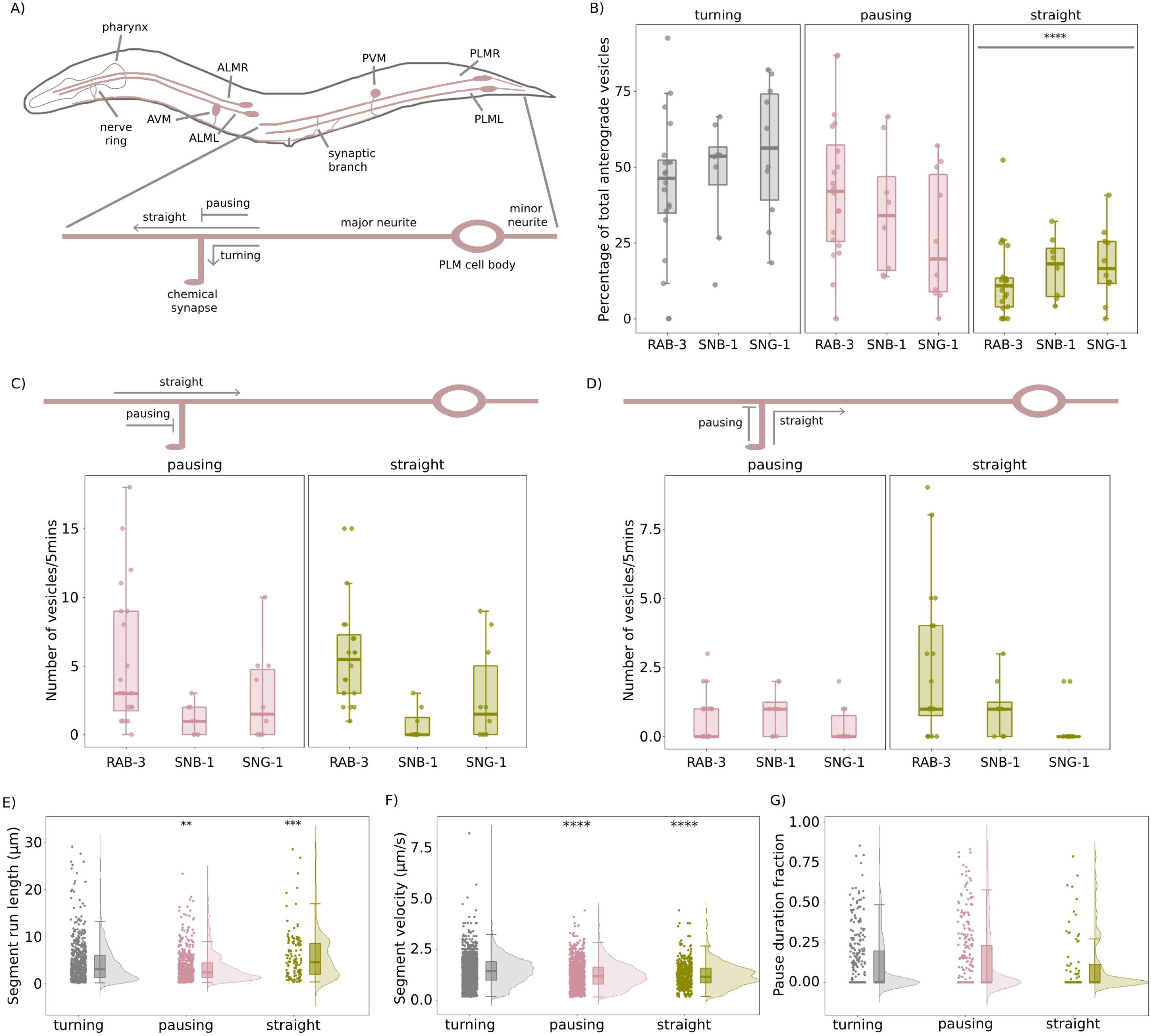
Characterisation of the behaviour of anterogradely-moving pre-SVs at the PLM branch point in wild type neurons. A) Schematic of the posterior lateral mechanosensory (PLM) neuron in *C. elegans*, representing the three behaviours exhibited by anterogradely-moving synaptic vesicles at the branch point. B) Proportion of ‘turning’, ‘pausing’, and ‘straight’ vesicles plotted as a percentage of total anterogradely-moving pre-SVs that reach the branch point in wild type (wt) animals. N(wt)=20 animals, n(wt-‘turning’)=303 vesicles, n(wt-‘pausing’)=309 vesicles, n(wt- ‘straight’)=76 vesicles. Wilcoxon rank sum test with continuity correction; ****p<0.0001; all comparisons are made to the ‘turning’ category. Individual data points are represented as solid circles. The summary statistics are depicted using a boxplot with whiskers. Upper whisker=minimum(maximum value in the distribution, Q3 + 1.5 * IQR) and lower whisker=maximum(minimum value in the distribution, Q1 – 1.5*IQR), where Q3 is the third quartile, Q1 is the first quartile, and IQR is the interquartile interval. C) Schematic of the *C. elegans* PLM neuron representing the two behaviours exhibited by retrogradely-moving vesicles from the asynaptic end at the PLM branch point. The plot represents total flux of pre-SVs RAB-3 (in *jsIs821*), SNB-1 (in *jsIs37*), and SNG-1 (in *jsIs219; tbIs222*) transported retrogradely from the asynaptic end of the PLM neuron over a period of 5 min. N(RAB-3)=20 animals, n(RAB-3-‘pausing’)=110 vesicles, n(RAB-3- ‘straight’)=118 vesicles. N(SNB-1)=8 animals, n(SNB-1-‘pausing’)=9 vesicles, n(SNB-1- ‘straight’)=6 vesicles. N(SNG-1)=10 animals, n(SNG-1-‘pausing’)=27 vesicles, n(SNG-1- ‘straight’)=28 vesicles. Wilcoxon rank sum test with continuity correction; all comparisons made to the ‘pausing’ category are not statistically significant. D) Schematic of the *C. elegans* PLM neuron representing the two behaviours exhibited by retrogradely-moving vesicles from the synaptic branch at the PLM branch point. The plot represents total flux of pre-SVs RAB-3 (in *jsIs821*), SNB-1 (in *jsIs37*), and SNG-1 (in *jsIs219; tbIs222*) transported retrogradely from the synaptic branch of the PLM neuron over a period of 5 min. N(RAB-3)=20 animals, n88(RAB-3-‘pausing’)=13 vesicles, n(RAB-3- ‘straight’)=49 vesicles. N(SNB-1)=8 animals, n(SNB-1-‘pausing’)=7 vesicles, n(SNB-1- ‘straight’)=8 vesicles. N(SNG-1)=10 animals, n(SNG-1-‘pausing’)=4 vesicles, n(SNG-1- ‘straight’)=4 vesicles. Wilcoxon rank sum test with continuity correction; all comparisons made to the ‘pausing’ category are non-significant. For all subsequent plots shown in the figure, individual data points are represented as solid circles. The distribution of the data is represented as a half-violin plot, while summary statistics are depicted using a boxplot with whiskers. Upper whisker=minimum(maximum value in the distribution, Q3 + 1.5 * IQR) and lower whisker=maximum(minimum value in the distribution, Q1 – 1.5*IQR), where Q3 is the third quartile, Q1 is the first quartile, and IQR is the interquartile interval. E) Segment run lengths (μm) of anterogradely-moving synaptic vesicles compared between turning, pausing, and straight vesicles in wild type. N(wild type)=20 animals, n(turning)=303 vesicles and 722 segments, n(pausing)=309 vesicles and 610 segments, n(straight)=76 vesicles and 191 segments. Wilcoxon rank sum test with continuity correction; **p<0.01, ***p<0.001; all comparisons are made to the ‘turning’ category. F) Segment velocities (μm/s) of anterogradely-moving synaptic vesicles compared between turning, pausing, and straight vesicles in wild type. N(wild type)=20 animals, n(turning)=303 vesicles and 2919 segments, n(pausing)=309 vesicles and 1681 segments, n(straight)=76 vesicles and 855 segments. Wilcoxon rank sum test with continuity correction; ****p<0.0001; all comparisons are made to the ‘turning’ category. G) Pause duration fraction of anterogradely-moving pre-SVs compared between turning, pausing, and straight vesicles in wild type. N(wild type)=20 animals, n(turning)=303 vesicles, n(pausing)=309 vesicles, n(straight)=76 vesicles. Welch’s Two Sample *t*-test; all comparisons made to the ‘turning’ category are non-significant.

## Results

### Synaptic vesicle precursors preferentially enter the synaptic branch or pause at the branch point

pre-SVs in the *C. elegans* PLM neuronal process (Fig. 1A) are observed by fluorescently-tagged synaptic vesicle marker proteins, such as RAB-3 (Rab GTPase), SNB-1 (Synaptobrevin-1), and SNG-1 (Synaptogyrin-1), over a period of 5 min using time-lapse fluorescence imaging. pre-SVs transported away from the PLM cell body towards the termini are termed ‘anterogradely-moving pre-SVs’, while pre-SVs transported back towards the cell body are termed ‘retrogradely-moving pre-SVs’. Anterogradely-moving pre-SVs exhibit three distinct behaviours at the PLM branch point: i) turning smoothly (without pauses) into the synaptic branch, ii) pausing at the branch point, and iii) going straight towards the asynaptic end (Figs. 1A, S1A). Anterogradely-moving pre-SVs that reach the PLM branch point predominantly turn into the synaptic branch (∼40–45%), or pause at the branch point (∼40–45%), while the proportion transported straight towards the asynaptic end is significantly lower (∼10%) (Fig. 1B, Supplementary movie 1). Retrogradely-moving pre-SVs at the branch point come from either the asynaptic end or the synaptic branch, which either pause at the branch point or continue straight along the main process back to the cell body (Fig. S1B). These pre-SVs exhibit equal preference for pausing at the branch point and going straight back towards the cell body (Figs. 1C, 1D). Additionally, the retrograde flux of pre-SVs is much lower than the anterograde flux of pre-SVs at the branch point (Fig. S1D). There are no retrogradely-moving pre-SVs that enter the synaptic branch directly from the main process, suggesting that there are no microtubule tracks in the main process whose minus-ends terminate in the branch.

### Synaptic vesicles entering the branch have distinct motion properties from vesicles in the main process

Motion properties of moving vesicles, such as run length, velocity, and pause duration fraction can be influenced by the number of microtubule-dependent motors on the vesicle surface, posttranslational modifications (PTMs) and microtubule-associated proteins (MAPs) on the underlying microtubule tracks, and presence of crowded environments (Sabharwal and Koushika, 2019, Vasudevan and Koushika, 2020). We find that anterogradely-moving pre-SVs turning into the branch have higher segment run lengths and velocities than pre-SVs that pause at the branch point (Figs. 1E, 1F). pre-SVs that travel straight along the main process have higher segment run lengths but lower segment velocities compared to the pre-SVs that turn into the branch (Figs. 1E, 1F). Pause duration fraction, a measure of the fraction of time a vesicle spends paused along its entire trajectory, is not significantly different between the turning, pausing, and straight populations of anterogradely-moving pre-SVs (Fig. 1G). The difference in run lengths and anterograde velocities between these populations may be partly explained by differences in the number of anterograde motors that drive their transport. To test this, we examined the effect of perturbing the levels of the anterograde motor on pre-SV transport at the branch point.

### Reduction in levels of functional anterograde motor UNC-104/KIF1A impedes the ability of pre-SVs to enter the synaptic branch

UNC-104/KIF1A is necessary for the anterograde transport of synaptic vesicles along neuronal processes in *C. elegans* (Hall & Hedgecock, 1991). In loss-of-function mutants of *unc-104 (unc-104(e1265tb120)* (*unc-104(lf)*), pre-SVs exhibit significantly reduced transport as compared to that seen in wild type (Kumar et al., 2010). The UNC-104 motors encoded by this allele are reported to have cargo binding defects, and pre-SVs have fewer motors on their surface (Kumar et al., 2010). We examined pre-SV behaviour at the PLM branch point in *unc-104(lf)/+* heterozygous animals, which have one functional copy of UNC-104 and another that exhibits cargo binding defects (Kumar et al., 2010). In these mutants, anterogradely-moving pre-SVs predominantly pause at the branch point (∼80% of all anterogradely-moving pre-SVs, Fig. 2A), and the proportion of pre-SVs turning into the branch or going straight are equally low (∼10% each, Fig. 2A, Supplementary movie 2). Reduction in levels of functional UNC-104 does not perturb the distribution of acetylated microtubules at the branch point (Figs. S2B, S2C), thus its influence on pre-SV behaviour is likely independent from the microtubule geometry at the branch point. The total number of anterograde pre-SVs at the branch point in *unc-104(lf)/+* heterozygous animals is significantly lower than wild type (Fig. S2D).

**Figure 2:**
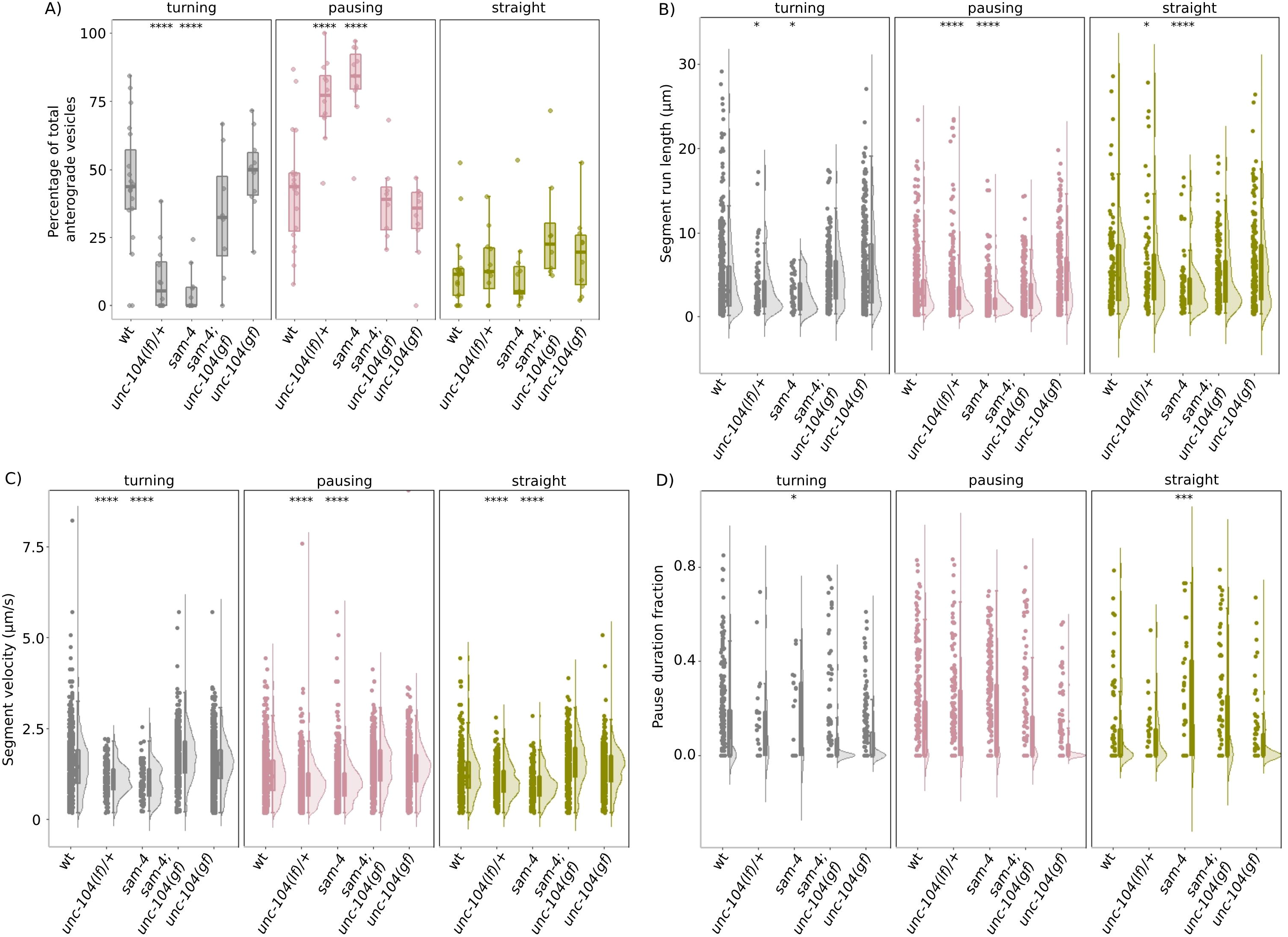
Reduction in levels of functional UNC-104 perturbs turning behaviour of anterograde pre-SVs. For the plots shown in this figure, individual data points are represented as solid circles. The distribution of the data is represented as a violin plot, while summary statistics are depicted using a boxplot with whiskers. Upper whisker=minimum(maximum value in the distribution, Q3 + 1.5 * IQR) and lower whisker=maximum(minimum value in the distribution, Q1 – 1.5*IQR), where Q3 is the third quartile, Q1 is the first quartile, and IQR is the interquartile interval. A) Proportion of ‘turning’, ‘pausing’, and ‘straight’ vesicles plotted as a percentage of total anterogradely-moving pre-SVs that reach the branch point in ‘wt’ (*jsIs821*), ‘*unc-104(lf)/+*’ [*unc-104(e1265tb120)/+;jsIs821*], ‘*sam-4*’ [*sam-4(js415);jsIs821*], ‘*sam-4 unc-104(gf)*’ [*sam-4(js415) unc-104(js1289);jsIs821*], and ‘*unc-104(gf)*’ [*unc-104(js1289);jsIs821*]. N(wt)=19 animals, n(wt-‘turning’)=306 vesicles, n(wt-‘pausing’)=265 vesicles, n(wt-‘straight’)=68 vesicles. N(*unc-104(lf)/+*)=12 animals, n(*unc-104(lf)/+*-‘turning’)=24 vesicles, n(*unc-104(lf)/+*-‘pausing’)=169 vesicles, n(*unc-104(lf)/+*-‘straight’)=35 vesicles. N(*sam- 4*)=11 animals, n(*sam-4*-‘turning’)=17 vesicles, n(*sam-4*-‘pausing’)=232 vesicles, n(*sam-4*- ‘straight’)=27 vesicles. N(*sam-4 unc-104(gf)*)=8 animals, n(*sam-4 unc-104(gf)*- ‘turning’)=151 vesicles, n(*sam-4 unc-104(gf)*-‘pausing’)=161 vesicles, n(*sam-4 unc-104(gf)*- ‘straight’)=86 vesicles. N(*unc-104(gf)*)=10 animals, n(*unc-104(gf)*-‘turning’)=192 vesicles, n(*unc-104(gf)*-‘pausing’)=142 vesicles, n(*unc-104(gf)*-‘straight’)=86 vesicles. Wilcoxon rank sum test with continuity correction; ****p<0.0001; all comparisons are made to the ‘wt’ category. B) Segment run lengths (μm) of anterogradely-moving pre-SVs compared between ‘wt’, ‘*unc-104(lf)/+*’, ‘*sam-4*’, ‘*sam-4 unc-104(gf)*’, and ‘*unc-104(gf)*’. N(wt)=19 animals, 639 vesicles, 1193 segments, N(*unc-104(lf)/+*)=12 animals, 228 vesicles, 561 segments, N(*sam- 4*)=11 animals, 259 vesicles, 684 segments, N(*sam-4 unc-104(gf)*)=8 animals, 398 vesicles, 817 segments, N(*unc-104(gf)*)=10 animals, 420 vesicles, 846 segments. Welch’s Two Sample *t*-test; *p<0.05, ****p<0.0001; all comparisons are made to the ‘wt’ category. C) Segment velocities (μm/s) of anterogradely-moving pre-SVs compared between ‘wt’, ‘*unc-104(lf)/+*’, ‘*sam-4*’, ‘*sam-4 unc-104(gf)*’, and ‘*unc-104(gf)*’. N(wt)=19 animals, 639 vesicles, 5059 segments. N(*unc-104(lf)/+*)=12 animals, 228 vesicles, 1779 segments, N(*sam- 4*)=11 animals, 259 vesicles, 1452 segments, N(*sam-4 unc-104(gf)*)=8 animals, 398 vesicles, 2359 segments, N(*unc-104(gf)*)=10 animals, 420 vesicles, 4058 segments. Welch’s Two Sample *t*-test; ****p<0.0001; all comparisons are made to the ‘wt’ category. D) Pause duration fraction of anterogradely-moving pre-SVs compared between ‘wt’, ‘*unc- 104(lf)/+*’, ‘*sam-4*’, ‘*sam-4 unc-104(gf)*’, and ‘*unc-104(gf)*’. N(wt)=19 animals, 639 vesicles. N(*unc-104(lf)/+*)=12 animals, 228 vesicles, N(*sam-4*)=11 animals, 259 vesicles, N(*sam-4 unc-104(gf)*)=8 animals, 398 vesicles, N(*unc-104(gf)*)=10 animals, 420 vesicles. Welch’s Two Sample *t*-test; comparisons made to the ‘wt’ category are non-significant.

It has been shown that a pre-SV-associated protein SAM-4, *C. elegans* homolog of vertebrate myrlysin, promotes UNC-104-mediated axonal transport of pre-SVs in neurons of *C. elegans* (Zheng et al., 2014, Niwa et al., 2017). We examined whether SAM-4 regulates pre-SV behaviour at the PLM branch point. Anterograde pre-SVs predominantly pause at the branch point in *sam-4(js415)* (null allele) mutant animals (∼90% of all anterogradely-moving pre-SVs) (Fig. 2A, Supplementary movie 3). The proportion of anterograde pre-SVs that turn into the branch in *sam-4* mutants is significantly reduced (∼5% of all anterograde pre-SVs), as is the total flux (Fig. S2D), while the proportion going straight is similar as compared to that seen in wild type (Fig. 2A).

The behaviour of anterograde pre-SVs in *sam-4* is phenotypically similar to that observed in *unc-104(lf)/+*. This suggests that *sam-4* and *unc-104* may function in the same genetic pathway to regulate pre-SV behaviour. A gain-of-function mutation in *unc-104*, *unc- 104(js1289)* (*unc-104(gf)*), suppresses the synaptic vesicle localisation defects observed in *sam-4* (Zheng et al., 2014). To determine whether *sam-4* and *unc-104* function in the same genetic pathway, we examined whether *unc-104(gf)* suppresses the pre-SV transport defects observed at the branch point in *sam-4*. We find that in *sam-4 unc-104(gf)*, the turning and pausing behaviour of anterograde pre-SVs is restored to wild type-like levels (Fig. 2A, Supplementary movie 4), while the proportion of pre-SVs going straight is not significantly altered (Fig. 2A), and the total flux of pre-SVs is significantly higher than wild type (Fig. S2D). The behaviour of anterograde pre-SVs at the branch point in *unc-104(gf)* mutants is similar to wild type (Fig. 2A, Supplementary movie 5).

Since the pre-SV localisation and transport defects observed in *sam-4* are suppressed by gain-of-function mutations in the motor domain of UNC-104, *sam-4* likely functions through UNC-104 to regulate pre-SV transport at the PLM branch point. SAM-4 is also reported to function as an upstream activator of UNC-104 through the release of its auto-inhibition via ARL-8 (Niwa et al., 2017). Since the turning, pausing, and straight populations of pre-SVs differ from each other in their motion properties, we investigated how a reduction in functional UNC-104 influences the motion properties of these populations at the branch point.

Anterograde pre-SVs in *unc-104(lf)/+* and *sam-4* show significantly reduced segment run lengths and velocities (Figs. 2B,2C) as compared to those seen in wild type, with *sam-4* showing more severe defects. pre-SVs also show a significant increase in the pause duration fraction in *sam-4* as compared to wild type (Fig. 2D). Since the turning population of pre-SVs is associated with high segment run lengths and segment velocities, the decrease in pre-SV run length and velocity observed in *unc-104(lf)/+* and *sam-4* is consistent with the loss of the turning population. The motion defects in *sam-4* are suppressed by *unc-104(gf)*, with pre-SVs in *sam-4 unc-104(gf)* exhibiting run lengths, velocities, and pause durations similar to those seen in wild type (Figs. 2B–2D). Anterogradely-moving pre-SVs in *unc-104(gf)* mutants do not exhibit any significant difference in motion properties compared to those seen in wild type (Figs. 2B–2D).

In summary, SAM-4/myrlysin functions to maintain the turning population of pre-SVs by regulating the activity of the anterograde motor UNC-104 on pre-SVs. Reduced activity of UNC-104, similar to the reduced levels of functional UNC-104, leads to the specific loss of the turning population and a concomitant increase in the pausing population of pre-SVs. Gain-of-function mutations in the motor domain of UNC-104 that increase the flux of the cargo-motor complex are sufficient to restore the turning behaviour of pre-SVs in *sam-4*, suggesting that the turning population of pre-SVs likely require increased activity of UNC- 104 as compared to the pausing and straight populations of pre-SVs.

### Increase in UNC-104 levels and activity influences vesicle behaviour at the branch point

Since we propose that the turning population of pre-SVs require higher levels of UNC-104 activity, we asked whether increasing the levels of UNC-104 in the neuron can increase the proportion of pre-SVs turning into the branch. Overexpression of UNC-104 in *C. elegans* neurons can function to increase UNC-104 activity, as it has been reported that overexpression of UNC-104 rescues the pre-SV mislocalization defects observed in *sam-4* (Zheng et al., 2014). We examined the behaviour of anterograde pre-SVs at the PLM branch point in the following conditions: i) overexpression of UNC-104 specifically in touch receptor neurons (henceforth termed as UNC-104 oe (PLM)), and ii) pan-neuronal overexpression of UNC-104 (henceforth termed as UNC-104 oe (pan neuronal)). Anterograde pre-SVs in UNC-104 oe (PLM) and UNC-104 oe (pan neuronal) show a slight but non-significant decrease in turning and pausing as compared to those seen in wild type, while the proportion going straight into the distal process is significantly increased (Fig. 3A, Supplementary movie 6 & 7). Increase in UNC-104 levels does not significantly alter the segment run lengths (Fig. 3B), segment velocities (Fig. 3C), and total flux of anterogradely-moving pre-SVs at the branch point (Fig. S2E). However, the pause duration fraction (Fig. 3D) of anterograde pre-SVs is significantly reduced in UNC-104 oe (PLM) and UNC-104 oe (pan neuronal) as compared to that seen in wild type. Overexpression of UNC-104 in the PLM neuron does not appear to grossly perturb its microtubule geometry at the branch point, as the distribution of acetylated microtubules at the branch point in UNC-104 oe (PLM) is similar to that in wild type (Fig. S2B, S2C).

**Figure 3:**
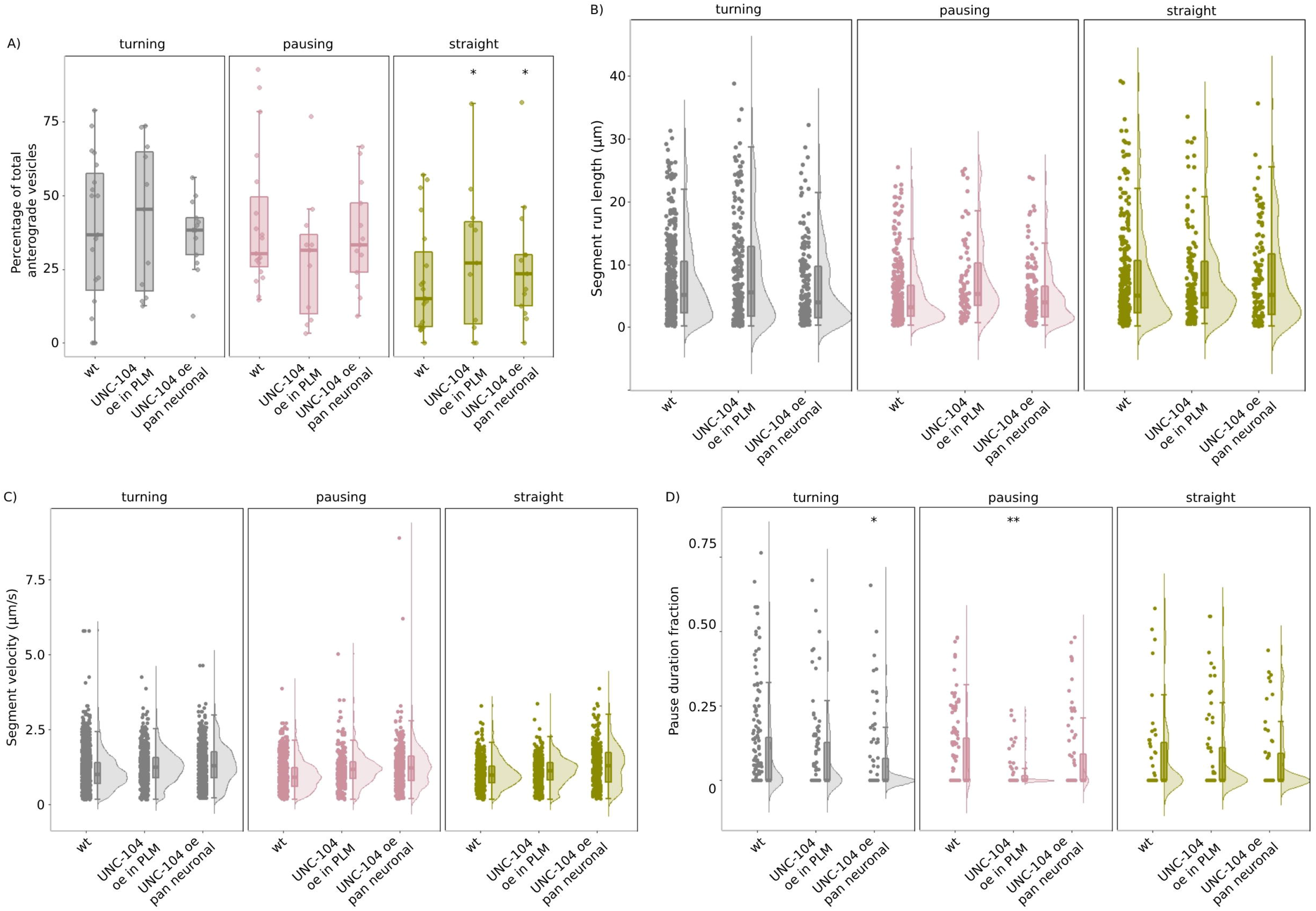
Increase in levels of UNC-104 drives more pre-SVs straight towards the asynaptic end. For the plots shown in this figure, individual data points are represented as solid circles. The distribution of the data is represented as a violin plot, while summary statistics are depicted using a boxplot with whiskers. Upper whisker=minimum(maximum value in the distribution, Q3 + 1.5 * IQR) and lower whisker=maximum(minimum value in the distribution, Q1 – 1.5*IQR), where Q3 is the third quartile, Q1 is the first quartile, and IQR is the interquartile interval. A) Proportion of ‘turning’, ‘pausing’, and ‘straight’ vesicles plotted as a percentage of total anterogradely-moving pre-SVs that reach the branch point in ‘wt’ (*tbIs227*), ‘UNC-104 oe in PLM’ (*jsIs1111;tbIs227*), ‘UNC-104 oe pan neuronal’ (*tbIs147;tbis227*). N(wt)=14 animals, n(wt-‘turning’)=152 vesicles, n(wt- ‘pausing’)=118 vesicles, n(wt- ‘straight’)=26 vesicles. N(UNC-104OE in PLM)=11 animals, n(UNC-104 oe in PLM-‘turning’)=94 vesicles, n(UNC-104 oe in PLM-‘pausing’)=55 vesicles, n(UNC-104 oe in PLM-‘straight’)=64 vesicles. N(UNC-104 oe pan neuronal)=13 animals, n(UNC-104 oe pan neuronal- ‘turning’)=82 vesicles, n(UNC-104 oe pan neuronal-‘pausing’)=76 vesicles, n(UNC-104 oe pan neuronal-‘straight’)=53 vesicles. Wilcoxon rank sum test with continuity correction; *p<0.05; all comparisons are made to ‘wt’. B) Segment run lengths (μm) of anterogradely-moving pre-SVs compared between wild type (wt), UNC-104 oe in PLM, and UNC-104 oe pan neuronal. N(wt)=14 animals, 296 vesicles, 1081 segments. N(UNC-104 oe in PLM)=11 animals, 213 vesicles, 432 segments. N(UNC- 104 oe pan neuronal)=13 animals, 211 vesicles, 429 segments. Welch’s Two Sample *t*-test; all comparisons are non-significant. C) Segment velocities (μm/s) of anterogradely-moving pre-SVs compared between wild type (wt), UNC-104 oe in PLM, and UNC-104 oe pan neuronal. N(wt)=14 animals, 296 vesicles, 5481 segments. N(UNC-104 oe in PLM)=11 animals, 213 vesicles, 2010 segments. N(UNC- 104 oe pan neuronal)=13 animals, 211 vesicles, 1630 segments. Welch’s Two Sample *t*-test; ***p<0.001; all comparisons are made to ‘wt’. D) Pause duration fraction of anterogradely-moving pre-SVs compared between wild type (wt), UNC-104 oe in PLM, and UNC-104 oe pan neuronal. N(wt)=14 animals, 296 vesicles. N(UNC-104 oe in PLM)=11 animals, 213 vesicles. N(UNC-104 oe pan neuronal)=13 animals, 211 vesicles. Welch’s Two Sample *t*-test; *p<0.05, **p<0.01; all comparisons are made to ‘wt’.

Providing higher levels of UNC-104 does not increase the turning population of pre-SVs, suggesting that the physiological levels of UNC-104 are sufficient to drive the maximum proportion of pre-SVs into the branch, and that turning behaviour is limited by other factors such as potentially the number of available microtubules tracks turning into the branch. Our observations also suggest that the physiological levels of active/functional UNC-104 are limiting for the straight population of anterograde pre-SVs. Increasing the levels of activated UNC-104 in the neuronal process drives more pre-SVs into the distal process compared to that in wild type, while the turning population is not significantly perturbed. It also reduces the propensity of pre-SVs to pause in their trajectory, thereby driving more pre-SVs straight into the distal process. Increase in UNC-104 levels appears to regulate the switch between the pausing and straight populations of anterogradely-moving pre-SVs, suggesting that in wild type neurons, the levels of UNC-104 are likely modulated to promote turning behaviour and limit the number of pre-SVs travelling to the asynaptic end.

## Discussion

Anterogradely-moving pre-SVs exhibit three distinct behaviours at the PLM branch point: i) turning into the synaptic branch, ii) pausing at the branch point, and iii) going straight into the distal process. This behaviour is observed across a heterogeneous population of pre-SVs (Fig. 1B), all of which depend on the anterograde motor UNC-104/KIF1A for transport out of the PLM cell body (Kumar et al., 2010, Maeder et al., 2014, Nadiminti et al., 2023). These distinct vesicle transport behaviours are found to differ in their sensitivity to the levels of functional UNC-104 in the neuronal process.

The population of pre-SVs turning into the branch are sensitive to low levels of UNC-104 (Fig. 2A) and unaffected by high levels of UNC-104 (Fig. 3A). This suggests that i) turning pre-SVs likely require more UNC-104 than pausing pre-SVs, ii) UNC-104 likely functions to limit the pausing population by promoting fast, processive anterograde transport with fewer pauses, iii) physiological levels of UNC-104 are not limiting for turning behaviour of vesicles. It is possible that anterograde pre-SVs already exhibit the maximum possible extent of turning that is allowed by the number of curved microtubules entering the branch. A reduction in levels/activity of UNC-104 does not significantly alter the proportion of pre-SVs going straight into the distal process (Fig. 2A). Since the transport of pre-SVs into the distal process is dependent on UNC-104, the insensitivity of the straight population to reduced levels of UNC-104 indicates that this proportion likely cannot go any lower (floor effect). This suggests that in wild type neurons, the proportion of pre-SVs going straight is at the lowest possible limit, given the underlying microtubule geometry at the branch point. Increasing the levels of active UNC-104 causes a significant increase in the proportion of pre-SVs going straight into the distal process (Fig. 3A). These observations collectively suggest that in, wild type neurons, the population of pre-SVs going straight can be limited in proportion due to the following reasons: i) vesicles going straight require higher levels of UNC-104 than available physiologically; thus, the frequency of such events is low, or ii) vesicles going straight are captured or halted at the branch point due to its crowded environment, and only a small proportion of vesicles are able to navigate or bypass the crowded geometry to enter the distal process.

## Acknowledgements and Funding

The authors gratefully acknowledge support from the Department of Atomic Energy, Government of India (DAE) grants 12-R&D-IMS-5.02-0202 and 1303/2/2019/R&DII/ DAE/2079, and the Howard Hughes Medical Institute (HHMI) International Early Career Scientist (IECS) grant 55007425 (to S.P.K.). Salary support and research costs were provided by funding from the PRISM project at the Institute of Mathematical Sciences, funded by the DAE, the Tata Institute of Fundamental Research (TIFR)-DAE (to A.V. and N.R.), CSIR (to S.P.K. and S.A.) and the Department of Biotechnology post- doctoral fellowship, Ministry of Science and Technology, India (to K.M.).

## Materials and Methods

### Maintenance of *C. elegans* strains

*C. elegans* strains were grown at 20°C on NGM plates seeded with *Escherichia coli* OP50 using standard methods of worm maintenance (Brenner, 1974). The 1-day adult animals used for imaging experiments come from three consecutive generations of no starvation, and clean, uncontaminated food. The strains used are listed in the Supplementary Table.

### Time-lapse image acquisition

Live animals were anaesthetised in 3mM tetramisole in M9 buffer on 5% agarose pads. Imaging was conducted on an Olympus IX83 inverted fluorescence microscope, integrated with the Yokogawa CSU-X1 spinning disc scan head from Perkin Elmer UltraVIEW, and the Hamamatsu ImagEM C9100-13/14 EMCCD Camera. Software used for image acquisition was Volocity 6.0 by Perkin Elmer. Images of the PLM branch point were acquired in the Green/Red channels at frame rates of 5 frames per second (fps) or 3 fps, using a UPLSAPO 100×, 1.4 N.A. oil immersion objective. Typically, the length of the PLM process imaged spans ∼70–80 μm, and the length of the PLM synaptic branch imaged spans ∼5–20 μm. Time-lapse movies were taken for durations of 5–10 min for all genotypes. Photobleaching of fluorescence signals at the branch point was conducted using the spot photobleaching function of the UltraVIEW PhotoKinesis accessory, with PK cycles=365, Step size=24, Spot period=75 ms, Spot cycles=13, Spot size=small. For animals imaged in the Green channel, 488 nm laser was used for photobleaching at a laser power of 10–20%.

### Kymograph generation for PLM branch point movies

All image panels used for representation and analysis of time-lapse movies were generated using FIJI (ImageJ v1.52p). Experimental kymographs were generated using the ‘MultipleKymograph’ plugin. Plugins were downloaded from the NIH website with the following links: http://www.rsbweb.nih.gov/ij/ and http://www.emblheidelberg.de/eamnet/html/bodykymograph.html. The official download page for the ‘MultipleKymograph’ plugin has been discontinued and a modified plugin called ‘KymoResliceWide’ is distributed at the following link: https://imagej.net/plugins/kymoreslicewide.

In the kymographs, cargo moving in the retrograde direction (towards the cell body), and anterograde direction (away from the cell body) appear as sloped lines, while stationary cargo appear as vertical lines. A cargo is counted as moving if it has been displaced by at least 3 pixels in successive time frames. For each movie of the PLM branch point, a ‘branch kymograph’ and ‘straight kymograph’ are generated as follows (Fig. S1C):

a. Branch kymograph is generated by tracing a curved segmented line along the neuron from the branch to the pre-branch main process
b. Straight kymograph is generated by tracing a straight segmented line from the post-branch to the pre-branch region along the main process

### Kymograph analysis for PLM branch point movies

Since the branch and straight kymographs both share the same pre-branch region and only differ from each other beyond the branch point, a comparison of vesicle trajectories in both kymographs helps us identify the branch point (Fig. S1C, S1D). Consequently, a vesicle trajectory that starts in the pre-branch region and crosses the branch point in the branch kymograph is categorised as a ‘turning vesicle’. A vesicle trajectory that starts in the pre-branch region and crosses the branch point in the straight kymograph is categorised as a ‘straight vesicle’. A vesicle trajectory that starts in the pre-branch region and stops at the branch point in both the branch and straight kymographs is categorised as a ‘pausing vesicle’. Annotated vesicle trajectories are then binned into different anterograde and retrograde categories based on their segments of origin and end and total counts of vesicles are calculated using custom-made in-house ImageJ macros.

### Analysis of motion properties

For the analysis of moving vesicle trajectories, the following motion properties were analysed: a) segment run lengths, b) segment velocities, c) pause duration fraction. Segment run lengths are defined as the distance moved by vesicles between consecutive pauses. Segment velocities of vesicles are defined as the distance covered per unit time between two consecutive pauses. Pause frequency is calculated as the total number of pauses exhibited by a moving vesicle divided by the total distance covered by the vesicle. Pause duration fraction is the fraction of the total time in a vesicle’s trajectory that it spends in pauses. Quantification of motion properties were done using custom-made in-house ImageJ macros The codes for the calculation of motion properties can be found on GitHub (https://github.com/amruta2612/Fijimacros).

### Immunostaining

*C. elegans* adult animals were fixed in 2% paraformaldehdye-methanol fixative (with added β-mercaptoethanol (βME)) and incubated at −80°C overnight. The fixed animals were washed sequentially in Borate-Triton buffer (with added βME) and then incubated at 4°C overnight in mouse anti-acetylated tubulin primary antibody (Ab) (Sigma T6793), at a dilution of 1:100 in 0.5% PBST. Following incubation in primary Ab, animals were washed in PBST and incubated in goat anti-mouse Alexa Fluor 488 secondary antibody (Invitrogen A11001), at a dilution of 1:100 in 0.5% PBST at 4°C overnight. Immunostained animals were mounted on glass slides using a mounting medium comprising 0.1 g N-propyl gallate, 3.5 ml glycerol, and 1.5 ml of 100 mM Tris pH 9.5 in 0.5% PBST.

### Statistical analysis

Shapiro–Wilk’s test was used on each sample set to test the normality of the distribution. Welch’s *t*-test was used to compare the means of distributions that passed the test of normality but differ from each other in their variance and sample sizes. Wilcoxon rank sum test (non-parametric) with continuity correction was used on data sets that did not pass the test of normality and wherein the distributions across different categories varied. All statistical tests were conducted in R (version 3.6.1).

## Supplementary figure legends

**Supplementary figure 1:**
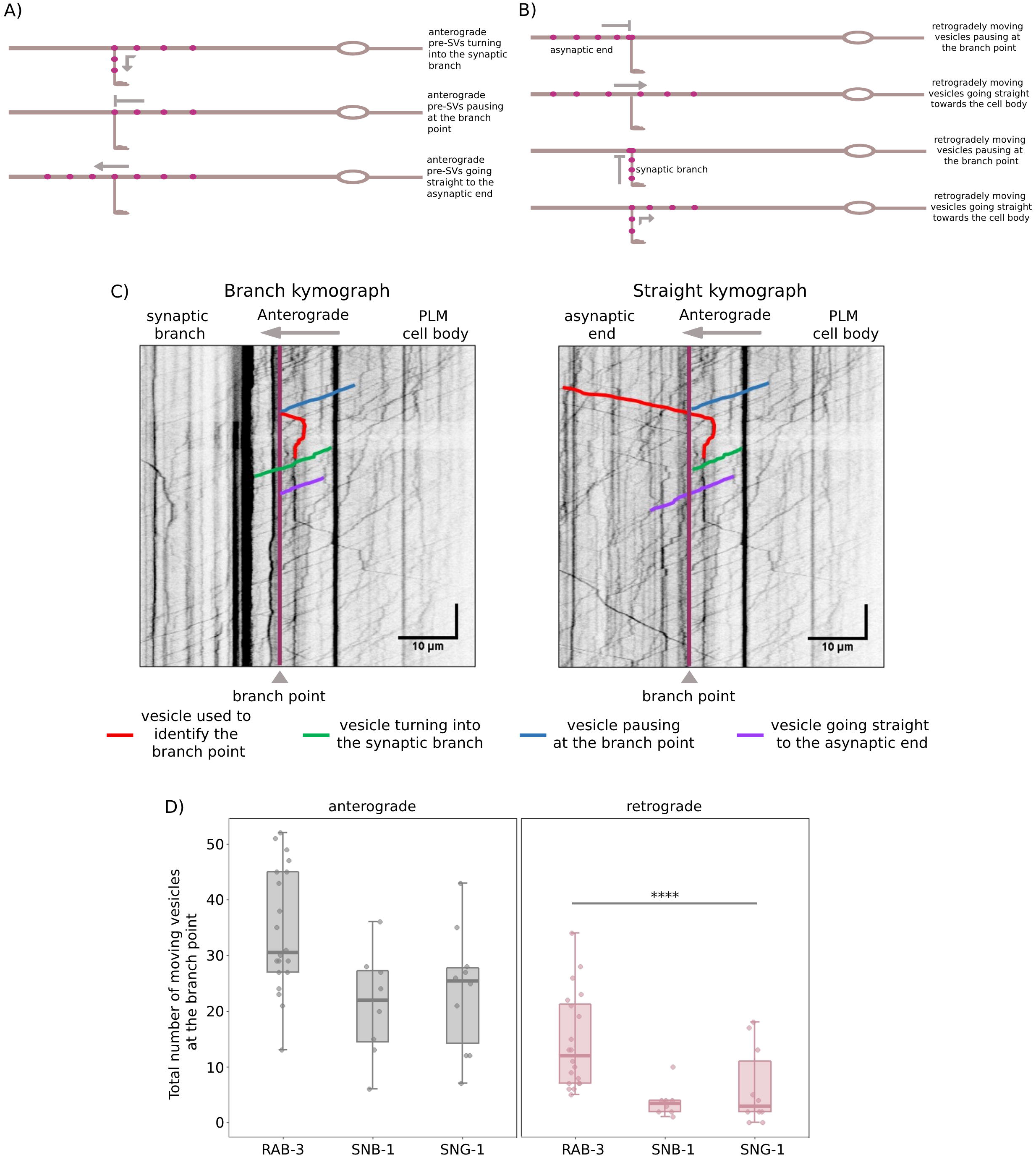
Analysis of pre-SV transport at the PLM branch point through kymographs. A) Schematic of the *C. elegans* PLM neuron representing the three behaviours exhibited by anterogradely-moving synaptic vesicles at the PLM branch point. B) Schematic of the PLM neuron representing the behaviours exhibited by retrogradely-moving vesicles from the asynaptic end and the synaptic branch at the PLM branch point. C) Representative kymographs depicting the identification and annotation of turning, pausing, and straight vesicles. X-axis scale bar=10 μm, Y-axis scale bar=5 s. D) Total number of pre-SVs transported in the anterograde and retrograde directions at the PLM branch point. N(‘RAB-3’)=20 animals, n(‘anterograde’)=639 vesicles, n(‘retrograde’)=263 vesicles. N(‘SNB-1’)=8 animals, n(‘anterograde’)=169 vesicles, n(‘retrograde’)=30 vesicles. N(‘SNG-1’)=10 animals, n(‘anterograde’)=236 vesicles, n(‘retrograde’)=63 vesicles. Wilcoxon rank sum test with continuity correction; all comparisons are made to the ‘anterograde’ category; ****p<0.0001.

**Supplementary figure 2:**
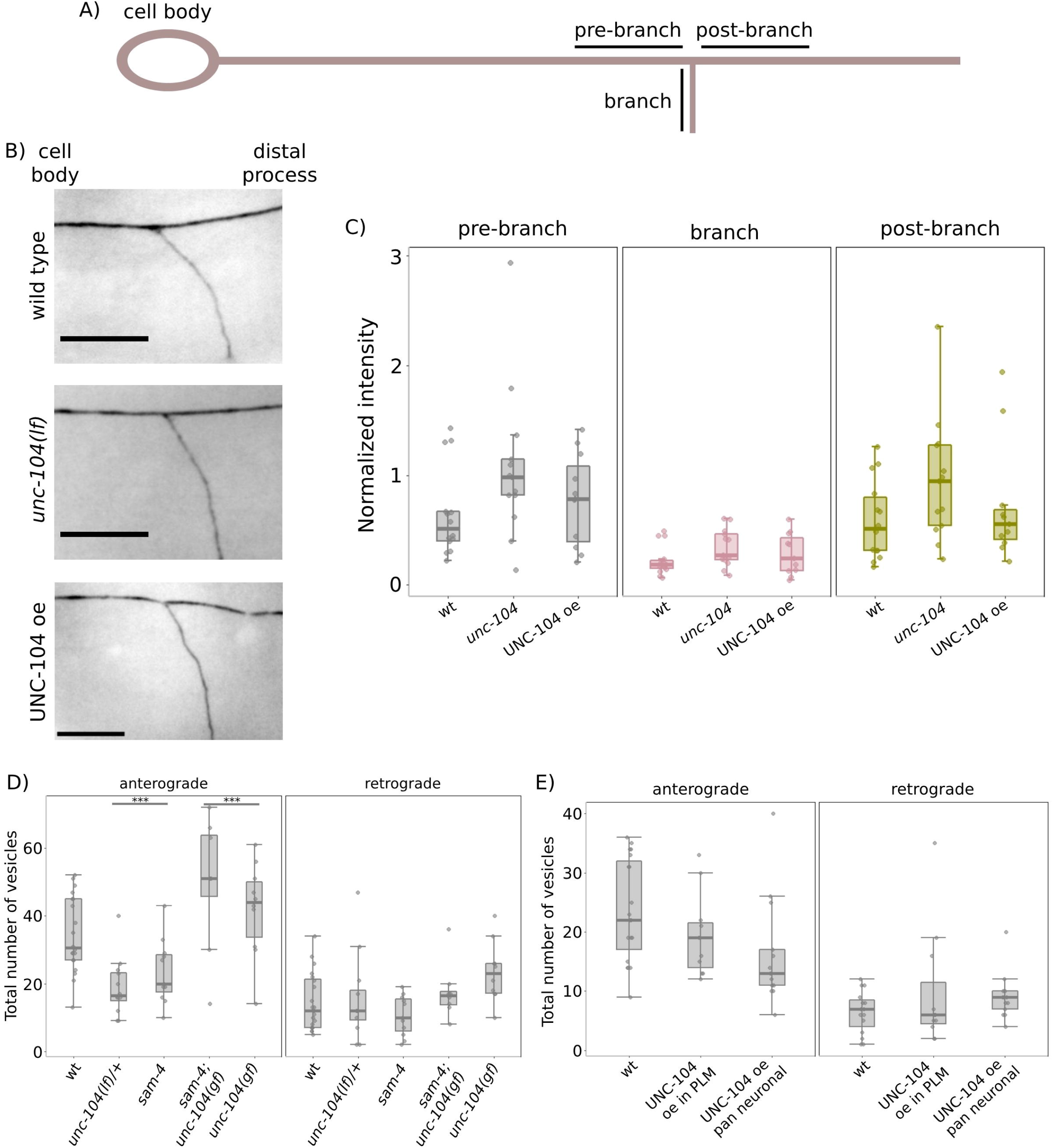
Changes in levels of UNC-104 do not affect the distribution of acetylated tubulin at the PLM branch point. A) Schematic of the posterior lateral microtubule (PLM) neuron, representing the ‘pre-branch’, ‘branch’, and ‘post-branch’ regions used for quantitation of signal from immunostaining. B) ‘wild type’ (N2), ‘*unc-104(lf)*’ [*unc-104(e1265tb120)*], ‘UNC-104 oe’ [*jsIs1111;tbIs227*]. Static representative images of acetylated tubulin staining at the PLM branch point. Images taken at 100×/1.4 N.A. scale bar=10 μm. C) Normalised intensity of acetylated tubulin staining compared between ‘wt’ (N2), ‘*unc- 104(lf)*’ [*unc-104(e1265tb120)*], ‘UNC-104 oe’ [*jsIs1111;tbIs227*]. N(wt)=14 animals, N(*unc-104*)=13 animals, N(UNC-104 oe)=11 animals. Wilcoxon rank sum test with continuity correction; all comparisons are made to the ‘wt’ category and are not statistically significant. D) Total number of pre-SVs transported in the anterograde and retrograde directions at the PLM branch point across different genotypes. N(‘wt’) = 20 animals, n(‘anterograde’)=639 vesicles, n(‘retrograde’)=263 vesicles. N(‘*unc-104(lf)/+*’) = 12 animals, n(‘anterograde’) = 228 vesicles, n(‘retrograde’) = 185 vesicles. N(‘*sam-4*’) = 11 animals, n(‘anterograde’) = 259 vesicles, n(‘retrograde’) = 117 vesicles. N(‘*sam-4 unc-104(gf)’*) = 8 animals, n(‘anterograde’) = 398 vesicles, n(‘retrograde’) = 141 vesicles. N(‘*unc-104(gf)’*) = 10 animals, n(‘anterograde’) = 420 vesicles, n(‘retrograde’) = 234 vesicles. Wilcoxon rank sum test with continuity correction; all comparisons are made to the ‘wt’ category; ***p<0.001. E) Total number of pre-SVs transported in the anterograde and retrograde directions at the PLM branch point across different genotypes. N(‘wt’) = 15 animals, n(‘anterograde’) = 338 vesicles, n(‘retrograde’) = 97 vesicles. N(‘UNC-104 oe in PLM’) = 11 animals, n(‘anterograde’) = 213 vesicles, n(‘retrograde’) = 107 vesicles. N(‘UNC-104 oe pan neuronal’) = 13 animals, n(‘anterograde’) = 211 vesicles, n(‘retrograde’) = 118 vesicles. Wilcoxon rank sum test with continuity correction; all comparisons are made to the ‘anterograde’ category.

**Supplementary Movie 1: pre-SVs turning into the synaptic branch in wild type *jsIs821* animals**

Anterogradely moving GFP::RAB-3 labelled pre-SVs shown turning into the PLM branch (marked by red arrow) in wild type, with the branch point marked by a green line. The PLM cell body is to the left and scale bar=5μm.

**Supplementary Movie 2: pre-SVs pausing at the branch point in *unc- 104(e1265tb120);jsIs821* mutants**

Anterogradely moving GFP::RAB-3 labelled pre-SVs (marked by red arrow) pausing at the branch point in *unc-104* mutants, with the branch point marked by a green line. The PLM cell body is to the left and scale bar=5μm.

**Supplementary Movie 3: pre-SVs pausing at the branch point in *sam-4(js415);jsIs821*** mutants

Anterogradely moving GFP::RAB-3 labelled pre-SVs (marked by red arrow) pausing at the branch point in *sam-4* mutants, with the branch point marked by a green line. The PLM cell body is to the left and scale bar=5μm.

**Supplementary Movie 4: pre-SVs turning into the synaptic branch in *sam-4(js415) unc- 104(js1289); jsIs821* animals**

Anterogradely moving GFP::RAB-3 labelled pre-SVs shown turning into the PLM branch (marked by red arrow), with the branch point marked by a green line. The PLM cell body is to the left and scale bar=5μm.

**Supplementary Movie 5: pre-SVs turning into the synaptic branch in *unc-104(js1289); jsIs821* animals**

Anterogradely moving GFP::RAB-3 labelled pre-SVs shown turning into the PLM branch (marked by red arrow), with the branch point marked by a green line. The PLM cell body is to the left and scale bar=5μm.

**Supplementary Movie 6: pre-SVs going straight into the distal process in *jsIs1111;tbIs227* animals**

Anterogradely moving mCherry::RAB-3 labelled pre-SVs shown going straight into the distal process (marked by red arrow), with the branch point marked by a green line. The PLM cell body is to the left and scale bar=5μm.

**Supplementary Movie 7: pre-SVs going straight into the distal process in *tbIs147;tbIs227* animals**

Anterogradely moving mCherry::RAB-3 labelled pre-SVs shown going straight into the distal process (marked by red arrow), with the branch point marked by a green line. The PLM cell body is to the left and scale bar=5μm.

