## Supplemental Table 1 for "Preferential transport of synaptic vesicles across neuronal branches is regulated by the levels of the anterograde motor UNC-104/KIF1A"

**Supplementary Table 1: List of strains**

| **S.No** | **Strain number** | **Genotype** | **Reference** |
| --- | --- | --- | --- |
| 1 | N2 | Bristol wild type | Brenner, 1974 |
| 2 | NM2689 | *jsIs821* [*mec-7*p::*gfp*::*rab-3*] | Bounoutas et al., 2009 |
| 3 | TT1884 | *tbIs227* [*mec-4*p::*mCherry*::*rab-3*] | Choudhary et al., 2017 |
| 4 | NM664 | *jsIs37* [*mec-7*p::*snb-1*::*gfp*] | Nonet, 1999 |
| 5 |  | *jsIs219* [*snb-1*p*::snb-1::gfp*]*;tbIs222* [*mec4*p*::mCherry*] | This study |
| 6 |  | *unc-104(e1265tb120); jsIs821* | Kumar et al., 2010 |
| 7 |  | *sam-4(js415);jsIs821* | Zheng et al., 2014 |
| 8 |  | *sam-4(js415) unc-104(js1289);jsIs821* | Zheng et al., 2014 |
| 9 |  | *unc-104(js1289);jsIs821* | Zheng et al., 2014 |
| 10 |  | *jsIs1111* [*mec4*p*:unc-104::gfp*] *;tbIs227* | Zheng et al., 2014, this study |
| 11 |  | *tbIs147* [*unc-104*p*:unc-104::gfp* in *unc-104(e1265)*] *;tbIs227* | Kumar et al., 2010, Sood et al., 2018 |
