## Supplementary figures and images for "Preferential transport of synaptic vesicles across neuronal branches is regulated by the levels of the anterograde motor UNC-104/KIF1A"

### Supplementary movie 1

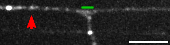

### Supplementary movie 2

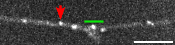

### Supplementary movie 3

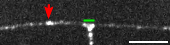

### Supplementary movie 4

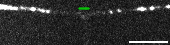

### Supplementary movie 5

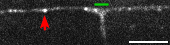

### Supplementary movie 6

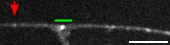

### Supplementary movie 7

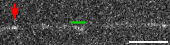
